# The electrophilic metabolite of kynurenine, kynurenine-CKA, targets C151 in Keap1 to derepress Nrf2

**DOI:** 10.1101/2025.11.18.689077

**Authors:** Jialin Feng, Mara Carreño, Hannah Jung, Sharadha Dayalan Naidu, Nicole Arroyo-Diaz, Abel D. Ang, Bhargavi Kulkarni, Dorothy Kisielewski, Takafumi Suzuki, Masayuki Yamamoto, John D. Hayes, Tadashi Honda, Beatriz Leon-Ruiz, Aimee L. Eggler, Dario A. Vitturi, Albena T. Dinkova-Kostova

**Author notes:** Co-corresponding authors: Albena T. Dinkova-Kostova, Jacqui Wood Cancer Centre, Division of Cancer Research, Ninewells Hospital and Medical School, James Arrott Drive, Dundee DD1 9SY, United Kingdom. and Dario A. Vitturi, Department of Pathology, School of Medicine, The University of Alabama at Birmingham, Birmingham, AL, USA. Co-first authors (Jialin Feng, Mara Carreño).

## Abstract

The Kelch-like ECH-associated protein 1 / Nuclear factor-erythroid 2 p45-related factor 2 (Keap1/Nrf2) system responds to a wide array of structurally diverse small molecules, of both exogenous and endogenous origin, by inducing a robust cytoprotective program that allows adaptation during oxidative, metabolic and inflammatory stress. Here, we report that exposure to the tryptophan metabolite kynurenine and its electrophilic derivative kynurenine-carboxyketoalkene (Kyn-CKA) leads to an increase in the abundance of transcription factor Nrf2 and induction of Nrf2-target genes, including NAD(P)H:quinone oxidoreductase 1 (NQO1), in murine and human cells. Additionally, both kynurenine and Kyn-CKA activate the aryl hydrocarbon receptor (AhR). Using cellular thermal shift assays, we found that Kyn-CKA increases the thermal stability of Keap1-mCherry fusion protein, but not free mCherry, indicating target engagement of Keap1, the principal repressor of Nrf2. The use of purified recombinant BTB domain of Keap1 and its C151S mutant counterpart revealed that Kyn-CKA reacts with wild-type, but not C151S mutant, Keap1-BTB, and at a much faster rate than with the small molecule thiol *N*-acetyl cysteine, demonstrating Kyn-CKA is targeted to react with C151 by the surrounding protein environment. In close agreement, Kyn-CKA increased the abundance of Nrf2 and expression of NQO1 in mouse embryonic fibroblast (MEF) cells expressing wild-type Keap1, but its inducer potency was greatly diminished in C151S-Keap1 mutant MEFs. Experiments in WT, AhR-knockout, and Nrf2-knockout primary murine bone marrow-derived macrophages showed that Nrf2 is required for the acute anti-inflammatory activity of Kyn-CKA, whereas AhR is dispensable. Together, these findings demonstrate that Kyn-CKA targets C151 in Keap1 to derepress Nrf2 and reveal that Nrf2, but not AhR, is a main contributor to the anti-inflammatory activity of Kyn-CKA in macrophages.

## 1. Introduction

In mammalian cells, the Kelch-like ECH-associated protein 1 / Nuclear factor-erythroid 2 p45-related factor 2 (Keap1/Nrf2) system is the main orchestrator of the cellular defence against environmental stress, most notably oxidative and inflammatory stress [1]. In this partnership, transcription factor Nrf2, which under homeostatic conditions is constitutively repressed by the activity of several ubiquitin ligase systems that promote its ubiquitination and subsequent proteasomal degradation, and Keap1, which is the substrate adaptor protein for the cullin 3 (Cul3)-really interesting new gene (RING) ubiquitin ligase complex, represents the principal negative regulator of Nrf2 [2]. During cellular stress, such as oxidative or electrophilic stress, Keap1 loses its repressor activity, and consequently *de novo* synthesized Nrf2 accumulates in the nucleus. This enables Nrf2 to heterodimerize with a small Maf protein and bind to the regulatory regions of genes that contain antioxidant/electrophile-responsive element (ARE/EpRE; i.e., 5ʹ-TGAC/GNNNGC-3ʹ) sequences, inducing a cytoprotective transcriptional program, which restores redox homeostasis. Prominent among the transcriptional targets of Nrf2 are genes encoding antioxidant enzymes, such as NAD(P)H:quinone oxidoreductase 1 (NQO1) and both the catalytic (GCLC) and the regulatory (GCLM) subunits of glutamate-cysteine ligase, which together catalyze the rate-limiting step in the biosynthesis of glutathione, the principal intracellular antioxidant. Additionally, Nrf2 activation has anti-inflammatory effects in cells, mice and humans, by suppressing the expression of pro-inflammatory genes, and thus plays a critical role in the resolution of inflammation [3–5].

Nrf2 can be activated pharmacologically by small molecules, the majority of which are electrophiles and oxidants that modify specific cysteine-based sensors in Keap1 [6,7]. Being in proximity to basic amino acids [8], C151 located in the Broad complex, Tramtrack and Bric-à-Brac (BTB) homodimerization domain of Keap1 (residues 51-180 [2]) is particularly reactive at physiological pH (p*K*_a_=6.9) [9], and recent work has revealed that, through mostly hydrophobic interactions, the reaction of C151 with structurally diverse Nrf2 activators is catalytic, with a key hydrogen bond orienting their electrophilic carbons within ∼3-5 Å of C151 for a catalytic proximity effect [9]. This is in full agreement with findings from mutagenesis studies showing that this cysteine is the main intracellular target for the most potent chemical class of inducers, the cyanoenones, irrespective of their molecular size and shape [10]. Notably, C151 in Keap1 is the target of the isothiocyanate sulforaphane, a classical Nrf2 activator that has been employed in ∼90 clinical trials, as well as for the two Nrf2 activators that are clinically in use: dimethyl fumarate, for relapsing remitting multiple sclerosis, and omaveloxolone, for Friedreich’s ataxia [11–13].

Numerous electrophilic Nrf2 activators belonging to several distinct chemical classes have been shown to suppress pro-inflammatory responses, with a linear correlation between their NQO1-inducer and inducible nitric acid synthase (iNOS)-inhibitory potencies spanning more than six orders of magnitude of concentrations [14,15]. Such striking correlation strongly suggests that, although the Keap1/Nrf2 system is not their only cellular target, it represents a significant contributor to the anti-inflammatory effects of these compounds. Most Nrf2 activators are xenobiotics, but it is becoming increasingly clear that several endogenously produced metabolites, particularly those that have electrophilic properties, can also react with cysteine-based sensors of Keap1 to activate Nrf2 [16].

Kynurenine is an endogenous metabolite derived from the essential amino acid tryptophan. It has recently been identified as a major metabolite within the tumour microenvironment, especially in aggressive cancers such as glioblastoma multiforme (GBM), where kynurenine is present at micromolar concentrations and promotes the establishment of an anti-inflammatory phenotype of tumour-associated macrophages, facilitating tumor growth [17]. Study of tumour-associated macrophages showed that the anti-inflammatory effect of kynurenine is mediated, in part, by activation of the aryl hydrocarbon receptor (AhR), a ligand-activated transcription factor that triggers expression of genes involved in numerous biological processes, including drug metabolism, haematopoiesis, angiogenesis and immunity [18]. Crosstalk between AhR and Nrf2 has been described, where the two transcription factors influence each other’s gene expression [19,20]. Notably, kynurenine and its metabolites, such as the electrophilic kynurenine-carboxyketoalkene (Kyn-CKA), have been demonstrated to activate Nrf2 in other pathologies, including sickle cell disease, attenuating inflammation [21]. Moreover, identification of the gene encoding the kynurenine-metabolising enzyme kynureninase as a gene transcriptionally upregulated by Nrf2 [22], provides a plausible negative feedback regulatory mechanism.

Because kynurenine is not electrophilic, whereas its metabolite Kyn-CKA is, we considered the possibility that Kyn-CKA is the actual Nrf2 activator. Using biochemical and cell-based assays, we found that Kyn-CKA reacts with C151 in the BTB domain of Keap1 and increases the thermostability of Keap1, indicating target engagement. Consequently, Nrf2 accumulates and induces transcription of ARE/EpRE-driven genes. We also show that in addition to activating Nrf2, Kyn-CKA inhibits the expression of genes encoding a panel of pro-inflammatory proteins in primary murine macrophages, in agreement with published work [21], and further reveal Nrf2, but not AhR, as one of the main contributors to the acute anti-inflammatory activity of Kyn-CKA in these cells.

## 2. Materials and methods

### 2.1. Materials

L-Kynurenine was purchased from Sigma (St. Louis, MO). CH-223191 (HY-12684) was from MedChemExpress LLC (Monmouth Junction, NJ), mouse M-CSF was from Prospec (Rehovot, Israel), Kyn-CKA [a mixture of two stereoisomers, (*E*)- and (*Z*)-(4-(2-aminophenyl)-4-oxobut-2-enoic acids; n.b., figure 3A shows only the structure of the *E* isomer] and Dean-CKA [(*E*)-4-oxo-4-phenylbut-2-enoic acid)] were custom-synthesized by LGC Standards (Manchester, NH). (±)-TBE-31 was synthesized as described previously [23]. All other chemicals were of analytical grade and obtained from Sigma unless specified. ELISA kits for MCP1 (DY479 and MJE00B) and IL6 (DY406) were obtained from R&D Systems (Minneapolis, MN). The AhR reporter assay kit (M06001) was obtained from INDIGO Biosciences (State College, PA). For this assay, L-kynurenine solutions were incubated with 35 mg of 3-mercaptopropyl-functionalized silica (SiliCycle, Quebec City, Canada) for 15 min at room temperature to remove potential Kyn-CKA traces, followed by 0.22-µm filtration.

### 2.2. Animals

Animal breeding and maintenance was in accordance with the regulations described in the UK Animals (Scientific Procedures) Act 1986, and with the approval by the Welfare and Ethical use of Animals Committee of the University of Dundee and the Institutional Animal Care and Use Committee of the University of Alabama at Birmingham (22775). Wild-type (WT), Nrf2-knockout (Nrf2-KO) and Keap1-knockdown (Keap1-KD) C57BL/6J mice [24] were bred and maintained at the WTB Resource Unit of the University of Dundee on a 12-h light/12-h dark cycle, 35% humidity with free access to water and pelleted RM1 diet (SDS Ltd., Witham, Essex, UK). In Nrf2-KO mice and derived cells, Nrf2 is transcriptionally inactive due to disruption of the *Nfe2l2* locus by an in-frame insertion of the *lacZ* coding sequence, which replaces part of exon 4 and all of exon 5 [25]. This knock-in retains the Neh2 Keap1-binding domain of Nrf2, resulting in Nrf2 turnover similar to the wild-type protein, but lacks the CNC-bZIP DNA binding and dimerization domains of the transcription factor. In Keap1-KD mice and derived cells, loxP sites inserted into the *Keap1* locus yielded a hypomorphic ‘knockdown’ allele with reduced *Keap1* expression even without Cre, resulting in constitutive Nrf2 stabilisation, increased nuclear Nrf2 levels, and elevated basal expression of Nrf2-target genes [26]. For the experiments shown in Figures 7-10, Nrf2-KO (B6.129X1-*Nfe2l2^tm1Ywk^*/J; RRID:IMSR_JAX:017009) and C57BL/6J (RRID:IMSR_JAX:000664) mice were purchased from the the Jackson Laboratory. *LysM AhR^-/-^* mice were generated in house by crossing LysM Cre (B6.129P2-Lyz2tm1(cre)Ifo/J; RRID:IMSR_JAX:004781) with *Ahr^flox/flox^* mice (Ahrtm3.1Bra/J; RRID:IMSR_JAX:006203).

### 2.3. Cell culture

All cultured cells were maintained in a humidified atmosphere of 95% air and 5% CO_2_ at 37°C, and were routinely tested to ensure that they were mycoplasma-free. MEF cells isolated from mice expressing wild-type or mutant Keap1 [27] and U2OS cells stably expressing Keap1-mCherry or free mCherry [28] were cultured in Dulbecco’s modified Eagle’s medium (DMEM), supplemented with 10% heat-inactivated FBS. ARPE-19 cells [28] were grown in DMEM, supplemented with 10% heat- and charcoal-inactivated FBS.

Mouse BMDM cells were obtained by *in vitro* differentiation of precursor cells isolated by flushing the tibia and femur with high-glucose DMEM containing 10% heat-inactivated FBS. Precursor cells were differentiated with either 20% L929 medium (for the experiments shown in Figures 1-3) or M-CSF (10 ng/mL, for the experiments shown in Figures 7-10) for 7 days in high-glucose DMEM, 10% heat-inactivated FBS and 1% penicillin/streptomycin. For the experiments shown in Figures 1-3, on day 7, adherent cells were detached with 5 mM EDTA in PBS (10 min, 37 °C), counted and replated in complete media, at the cell densities indicated in the legends. For the experiments shown in Figures 7-10, cells were seeded in 12-well plates (0.5 x 10^6^/well) in complete media and incubated at 37°C, 5% CO_2_. Experimental treatments were performed in the absence of heat-inactivated FBS for the indicated durations. For LPS treatments (*E. coli* O111:B4), cells were pre-treated with Kyn-CKA for 40 min prior to endotoxin addition.

**Figure 1.**
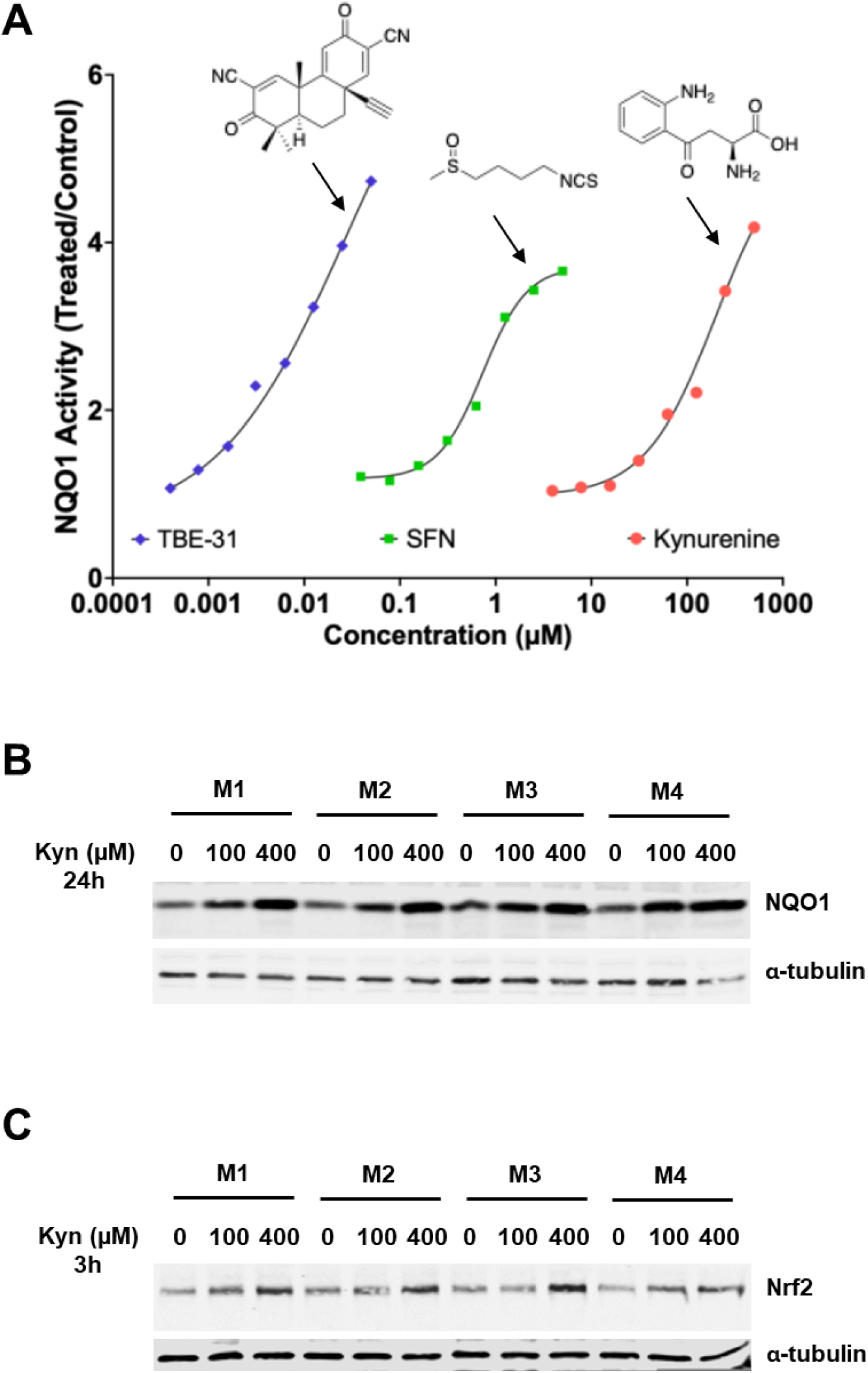
Kynurenine is an Nrf2 activator in BMDMs. **(A)** BMDMs were resuspended, seeded into 96-well plates (100,000 cells/well), and treated 24 h post-seeding with either vehicle [0.1% DMSO (v/v)] or increasing concentrations of kynurenine, TBE-31, or sulforaphane (SFN). After 48 h, the specific enzyme activity of NQO1 was quantified in cell lysates. Data are shown as fold change in NQO1 specific enzyme activity relative to vehicle control. Values represent means of 8 biological replicates. The standard deviation was <10% of each value. **(B,C)** BMDMs, derived from four WT C57BL/6J mice (M1-M4), were seeded into six-well plates (1 x 10^6^/well), allowed to adhere overnight, and subsequently treated with either vehicle or kynurenine (Kyn, 100 µM or 400 µM) for either 3 h or 24 h. Total cell lysates were prepared and the protein levels of NQO1 **(B)** were determined by immunoblotting 24 h post-treatment. The protein levels of Nrf2 **(C)** were determined by immunoblotting 3 h post-treatment. The levels of α-tubulin served as a loading control.

### 2.4. NQO1 enzyme activity assay

NQO1 inducer activity was determined using the "Prochaska" microtiter plate bioassay [29]. Cells were grown in 96-well plates for 24 h, and then exposed in 8 replicates to 2-fold serial dilutions of each compound or vehicle for a further 24 h or 48 h, as indicated in the figure legends. After washing with PBS (200 µL, twice), cell lysates were obtained by adding 75 µL digitonin (0.8 g/L in 2 mmol/L EDTA, pH 7.8) followed by incubation at room temperature for 10 min with shaking. Aliquots of cell lysates (20 µL) were transferred to parallel 96-well plates, and protein concentrations were measured by the bicinchoninic acid (BCA) assay (Thermo Fisher). The enzyme activity of NQO1 (with menadione as a substrate) was quantified using the remaining 55 µL of lysate. Inducer potency was expressed as a CD value, i.e., the Concentration that Doubles the NQO1 specific enzyme activity.

### 2.5. Immunoblotting

Following treatment, cells were washed twice with ice-cold PBS and lyzed in SDS lysis buffer [50 mM Tris pH 6.8, 10% glycerol (v/v), 2% SDS (w/v)]. The lysates were sonicated for 30 sec (using 5 sec on - 1 sec off cycles) at 20% amplitude, the protein concentrations were determined using the BCA assay (Thermo Fisher). Samples were adjusted to the same protein concentration, reduced by adding 2-mercaptoethanol to a final concentration of 5% (v/v), incubated for 30 min at room temperature and then heated at 95°C for 5 min before being subjected to electrophoresis. Bromophenol Blue [0.001% (w/v)] was added, and equal sample volumes containing equal amounts of total protein, were loaded onto either Tris-Glycine or 4–12% Bis-Tris NuPAGE gels (Thermo Fisher). Proteins were separated by electrophoresis using either Tris-Glycine or MOPS (Thermo Fisher) buffer, and transferred onto 0.45-μm premium nitrocellulose membranes (Amersham) by wet transfer (Bio-Rad).

Transfer efficiency was confirmed by membrane incubation with 0.1% (w/v) Ponceau S solution in 5% (v/v) acetic acid. Stain was then washed off by PBS-0.1% (v/v) Tween 20 (PBST), after which membranes were blocked in 5% (w/v) non-fat milk (Marvel) dissolved in PBST (milk-PBST) for 1 h at room temperature on an orbital shaker. The membranes were incubated with primary antibodies overnight at 4°C or for 1 h at room temperature for loading controls (e.g. GAPDH, α-tubulin or β-actin), washed with PBST (3 times, 10 min each), and incubated with fluorescently conjugated secondary antibodies (1:20000) (LI-COR) for 1 h at room temperature. The blots were then washed with PBST (3 times, 10 min each) at room temperature, and the proteins were visualized using the Odyssey CLx imager (LI-COR). The primary antibodies and their respective dilutions and sources were: Keap1 (1:4000, clone 144, Millipore MABS514); NQO1 (1:1000, clone D6H3A, Cell Signaling 62262); Nrf2 (1:1000, clone E5F1A, Cell Signaling 20733); GAPDH (1:30000, clone 1E6D9, Proteintech 60004-1-Ig); β-actin (1:10000, clone AC-15, Sigma-Aldrich A5441); and α-tubulin (1:10000, clone DM1A, Cell Signaling 3873). All antibodies were diluted in milk-PBST.

### 2.6. mRNA analysis

Cells were lysed and total RNA extracted either using PureLink™ RNA Mini Kit (Thermo Fisher) or TRIzol reagent (Invitrogen, USA; 0.4 mL per million cells) and quantified by UV spectroscopy at 260 nm or NanoDrop^TM^ One/OneC. Reverse transcription was performed using either PrimeScript™ RT Master Mix kit (Takara Bio) or iScript cDNA kit (Bio-Rad). Real-time PCR was carried out on Applied Biosystems QuantStudio™ 5 or 7 Real-Time PCR systems using TaqMan Fast Advanced Master Mix (Thermo Fisher) or PowerUp™ SYBR™ Green Master Mix (Thermo Fisher). Results were expressed as fold changes in gene expression over actin using the ΔΔCt method. The TaqMan™ Gene Expression Assay IDs (Thermo Fisher, Waltham, MA) were: Mm01253561 (for *Nqo1*); Mm00516005 (for Ho1); Mm01324400 (for *Gclm*); Mm00434228 (for *IL1b*); Mm00446190 (for *IL6*); Mm00441242 (for *Mcp1*); Mm00443252 (for *Tnf*); Mm004405021 (for *Nos2*); Mm00437762 (for *B2m*); Mm03024075 (for *Hprt1*); and 4352341E (for *Actb*). The sequences of the primers for the SYBR™ Green Real-Time PCR were:

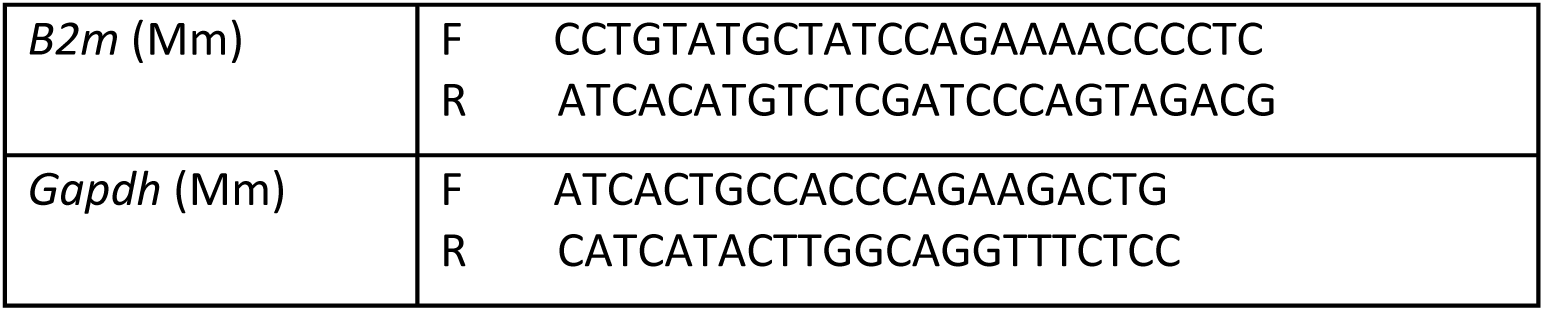
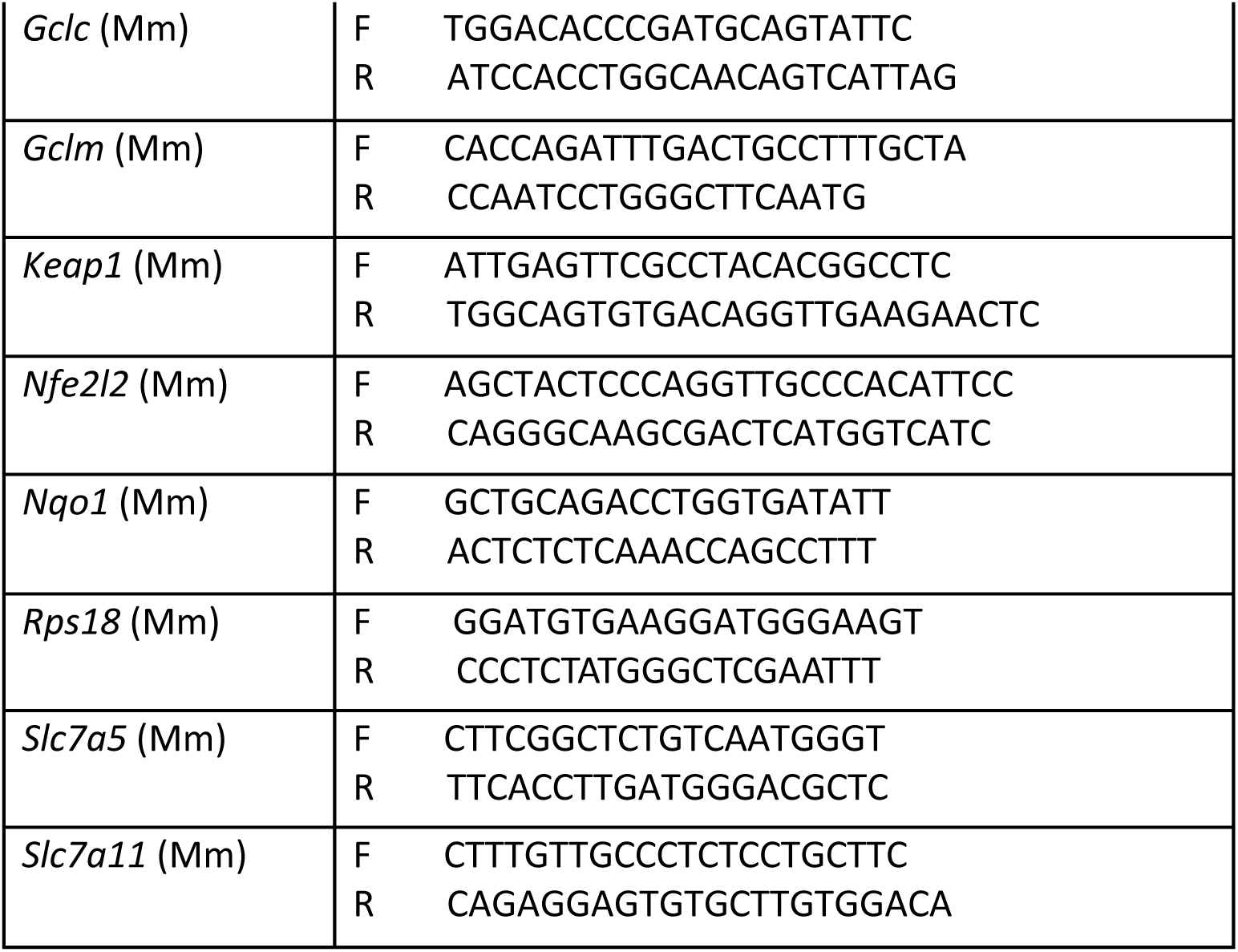

### 2.7. Kyn-CKA reactions with N-acetyl cysteine or BTB-C151

The WT Keap1 BTB domain protein and the BTB-C151S mutant protein were expressed and purified as previously described [9]. The proteins were reduced with 1 mM DTT at room temperature for 1 h and subsequently buffer exchanged into degassed 100 mM phosphate buffer pH 7.5 using Zeba 7 kDa MWCO spin desalting columns. To avoid rapid oxidation of C151, the protein was used within 30 min or stored under argon. Kyn-CKA was preincubated in pH 7.5 100 mM phosphate buffer for 2 h then reacted with *N*-acetyl cysteine, WT BTB, or BTB-C151S (or buffer control) in the same buffer. Reactions were initiated by adding Kyn-CKA to wells containing *N*-acetyl cysteine, WT BTB, BTB-C151S, or buffer as a control. Final concentrations were 30 µM Kyn-CKA and 30 µM BTB-C151 or BTB-C151S. Given the C151 p*K*_a_ of 6.9 [9], 24 µM C151 thiolate is available to react at pH 7.5. Correspondingly, to obtain a final reaction concentration of 24 µM thiolate *N*-acetyl cysteine, with a p*K*_a_ of 9.4, the final concentration of *N*-acetyl cysteine was 2.5 mM in the reactions. Absorbance spectra from 300 nm to 600 nm were collected on a BMG Labtech ClarioStar in kinetic scanning mode.

The change in absorbance at 410 nm over time (normalized by subtraction for the absorbance of BTB or *N*-acetyl cysteine alone) versus time was fitted to a one-phase decay, for the indicated data.

### 2.8. CETSA

CETSA was performed as described [28,30]. U2OS cells stably expressing Keap1-mCherry or mCherry were induced with doxycycline hyclate (0.5 μg/mL) for 48h. Cells were washed three times with ice-cold PBS and then collected by scraping into 1.0 mL PBS supplemented with protease inhibitor cocktail (Roche). Cell suspensions were snap-frozen in liquid nitrogen and stored at -70°C. Subsequently, cells were lyzed by six freeze-thaw cycles, and cell debris was removed by centrifugation at 9,600 x *g* at room temperature for 5 min. Clarified cell lysates were collected, and protein concentrations were measured using the BCA assay (Thermo Fisher). Samples were normalised, ensuring control and treated groups were adjusted to the same protein concentration, and then incubated with either Kyn-CKA or vehicle [0.1% MeOH (v/v)] for 1 h at 37°C. The incubation mixtures were aliquoted (100 μL) and heated for 3 min at the indicated temperatures (i.e. from 37°C to 70°C, in 3°C increments), and the aggregated material was removed by centrifugation at 17,000 x *g* at 4°C for 40 min. The fluorescence in the soluble fraction was measured with 580 nm excitation, 590 nm cut-off and 615 nm emission wavelengths.

For ITDRF-CETSA, cell lysates were aliquoted (100 μL) and incubated with a range of compound concentrations or vehicle [0.9% MeOH (v/v)] for 1 h at 37°C to allow compound-target interaction. Following incubation, samples were subjected to heat treatment at 64°C for 3 min. The aggregated material was removed by centrifugation and the fluorescence in the soluble fraction was measured as described above.

### 2.9. AhR reporter assay

The assay was performed following the manufacturer’s instructions. Briefly, 200 µL of mouse AhR reporter cells expressing luciferase were seeded in 96-well plates and incubated at 37°C and 5% CO_2_. After 6 h, the culture media was replaced with treatment media.

Reporter cells were treated with variable concentrations of L-Kynurenine, Kyn-CKA or Dean-Kyn-CKA (0 to 30 µM) for 24 h. Alternatively, they were pre-incubated with 10 µM CH223191 (AhR inhibitor) for 1 h, followed by treatment with 10 µM L-Kynurenine or Kyn-CKA, in the presence of 10 µM AhR inhibitor. After 24 h, the media were discarded, and Luciferase Detection Reagent was added followed by luminescence measurement using Spectramax iD5 (Molecular Devices) plate reader.

### 2.10. Quantification and statistical analysis

All quantitative data are represented graphically as mean values ± 1 standard deviation (SD) or standard error of means (SEM), as indicated in the figure legends. Student’s t-test (for two-group comparisons) or one-way ANOVA (for multiple comparisons) and either Tukey’s or Dunnett’s post-test (indicated in the figure legends) were used to test for statistical significance.

## 3. Results

### 3.1. Kynurenine induces Nrf2-driven transcription in murine bone marrow-derived macrophages

Our previous high-resolution proteomic analysis showed that the Keap1/Nrf2 system is fully functional in primary murine bone marrow-derived macrophages (BMDMs), and its genetic or pharmacologic activation suppresses their pro-inflammatory responses to lipopolysaccharide (LPS) whilst supporting their redox metabolism and mitochondrial adaptation to inflammatory stress [31]. Thus, we employed this system to explore the possibility that kynurenine activates Nrf2 in a physiologically relevant cellular model using a quantitative bioassay that measures the enzyme activity of the classical Nrf2 target NQO1 [32]. Treatment with kynurenine for 48 h led to a concentration-dependent increase in NQO1 specific enzyme activity (**Figure 1A**). However, the concentration required to double the NQO1 enzyme activity (CD value) was higher (CD=100 µM) compared to two other well-established NQO1 inducers, the tricyclic cyanoenone TBE-31 (CD=0.03 µM) and the isothiocyanate sulforaphane (SFN, CD=0.6 µM), indicating that kynurenine is a relatively low potency NQO1 inducer. The concentration-dependent induction of NQO1 by kynurenine was further confirmed by immunoblotting of lysates of cells that had been treated with kynurenine for 24 h (**Figure 1B**); this type of analysis also showed that the 3 h-kynurenine treatment increased the abundance of Nrf2 protein (**Figure 1C**).

To examine whether the kynurenine-mediated upregulation of NQO1 (and other Nrf2-transcriptional targets) is dependent on Nrf2, RT-qPCR analysis was performed on BMDMs isolated from wild-type (WT) or *Nfe2l2*^⁻/⁻^ (i.e., Nrf2-KO) mice. In cells from Nrf2-KO mice, Nrf2 is transcriptionally inactive [25]. BMDMs of the two genotypes were treated with vehicle, 20 µM, or 200 µM kynurenine for 18 h, and the mRNA levels for *Nfe2l2*, *Keap1*, and the Nrf2 target genes *Nqo1*, *Gclc*, *Gclm*, *Cd36* and *Slc7a11* were quantified. Notably, the *Nfe2l2* mRNA levels were substantially lower in vehicle-treated Nrf2-KO BMDMs (**Figure 2A**). Although to a much smaller extent, the *Keap1* mRNA levels were also lower in vehicle-treated Nrf2-KO BMDMs in comparison with their WT counterparts (**Figure 2B**).

**Figure 2.**
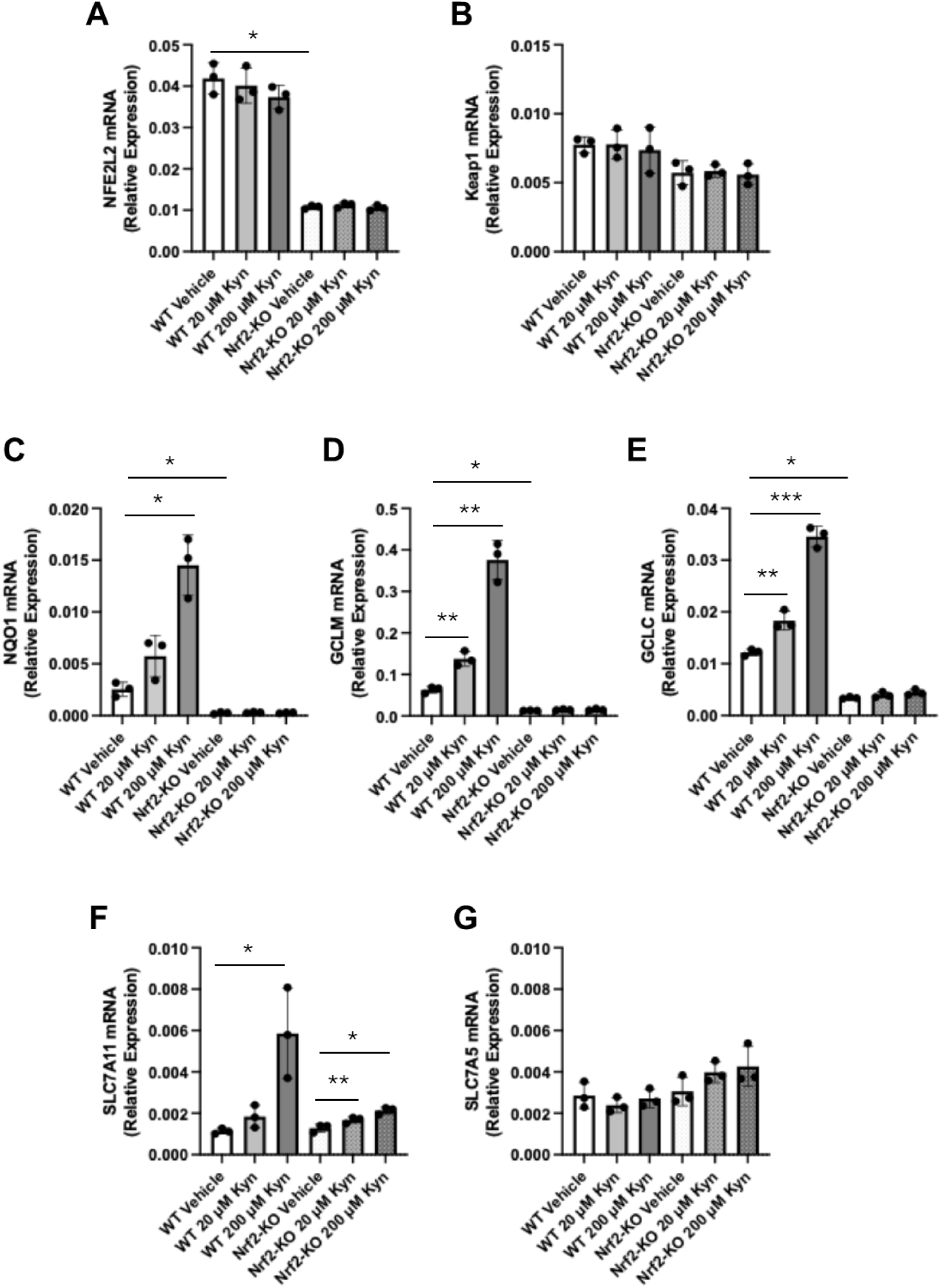
Kynurenine induces Nrf2-driven transcription in BMDMs. WT and Nrf2-KO BMDMs seeded into six-well plates (1 x 10^6^/well), allowed to adhere overnight, and then treated for 18 h with vehicle or kynurenine (Kyn; 20 µM or 200 µM). The levels of mRNA for Nrf2 (gene name *Nfe2l2*) **(A)**, *Keap1* **(B)**, and the Nrf2 transcriptional targets *Nqo1* **(C)**, *Gclm* **(D)**, *Gclc* **(E)**, *Slc7a11* **(F)**, and *Slc7a5* **(G)** were determined by quantitative RT-PCR after normalisation to the housekeeping control *Rps18* (or *B2m* for *Keap1*). Bars represent means ± SEM (n=3 biologically independent replicates per condition), with individual data points shown.*, p<0.05; **, p<0.01; ***, p<0.001; ****, p<0.0001.

In WT BMDMs, kynurenine caused a robust, concentration-dependent increase in the mRNA levels of the Nrf2-target genes, with *Nqo1* and *Gclm* showing greater than five-fold upregulation in cells treated with 200 µM kynurenine (**Figure 2C-E**). In contrast, in Nrf2-KO BMDMs, the basal and the inducible expression of *Nqo1*, *Gclc*, *Gclm*, and *Cd36* was strongly suppressed, or absent, demonstrating clear Nrf2 dependence. One exception was *Slc7a11* (**Figure 2F**), whose basal levels were similar between WT and Nrf2-KO cells, and its expression remained partially inducible in Nrf2-KO BMDMs, albeit to a much smaller (1.7-fold in Nrf2-KO vs. 5.2-fold in WT BMDMs treated with 200 µM kynurenine) magnitude. This result suggests that the expression of *Slc7a11* involves additional transcriptional inputs, such as ATF4, which has also been shown to be activated by kynurenine [33]. Importantly, the expression of *Slc7a5* was unaffected by neither genotype nor kynurenine treatment (**Figure 2G**), indicating that the transport capacity of large neutral amino acids (including tryptophan and kynurenine) is constitutive and Nrf2-independent in this system. Together with previously published work [21,33], these results establish that the endogenous metabolite kynurenine is an inducer of a broad transcriptional program in macrophages via a mechanism that is dependent on Nrf2.

### 3.2. Kynurenine-CKA is a Nrf2-dependent inducer of NQO1

High levels of kynurenine in cells, and *in vivo*, lead to the generation of the electrophilic metabolite kynurenine-carboxyketoalkene (Kyn-CKA), through spontaneous deamination [21]. Unlike kynurenine, which is not electrophilic, Kyn-CKA contains an α,β-unsaturated carbonyl, a Michael acceptor moiety, which is a characteristic feature of many compounds that inhibit Keap1, block ongoing proteasomal degradation of Nrf2, and induce NQO1 activity [34,35]. Treatment with Kyn-CKA was previously shown to induce Nrf2-dependent gene expression in mice *in vivo* as well as in the murine hepatocyte cell line AML-12 and MEF cells derived from wild-type, but not Nrf2-knockout mice [21]. These data, together with the electrophilic properties of Kyn-CKA and the high concentrations of kynurenine that were required to induce NQO1, led us to hypothesize that Kyn-CKA, and not kynurenine itself, was the actual inducer.

To test this hypothesis, we performed a small structure-activity relation study comparing the NQO1 inducer activity of kynurenine and Kyn-CKA, in which we also included the following: (i) Dean-Kyn-CKA, the deaminated derivative of Kyn-CKA, which lacks the *ortho*-aminophenyl group (NH_2_), and so prevents intramolecular cyclisation that would otherwise lead to loss of electrophilicity; (ii) Red-Kyn-CKA, a non-electrophilic form of Kyn-CKA in which the α,β-unsaturated carbonyl is reduced; (iii) kynurenic acid (KynA), an endogenous biologically active non-electrophilic metabolite of kynurenine. For this comparison, we used the human ARPE-19 cell line, which we have previously shown to respond robustly to compounds that increase Nrf2 activity [36]. Both Kyn-CKA and its deaminated derivative (Dean-Kyn-CKA) produced a clear concentration-dependent induction of NQO1 (**Figure 3A**), with Dean-Kyn-CKA displaying higher potency (CD = 10 µM) than Kyn-CKA (CD = 30 µM). As expected, kynurenine also induced NQO1 in a concentration-dependent manner, but with much lower potency (CD = 400 µM). KynA elicited only a very slight (1.2-fold) increase in NQO1 activity at the highest concentrations tested, and the reduced, non-electrophilic analog (Red-Kyn-CKA) was inactive. Together, these data define a clear structure-activity relationship where the electrophilicity of Kyn-CKA is essential for robust NQO1 induction, and also further strengthen the notion that Kyn-CKA rather than kynurenine itself, is the actual inducer. In agreement with these results in ARPE-19 cells, Kyn-CKA was also much more potent than kynurenine in inducing NQO1 in BMDMs: the NQO1 mRNA level was increased by 5.7-fold after treatment with 30 µM Kyn-CKA (**Figure 3B**), whereas this level of induction was achieved by 200 µM kynurenine (**Figure 2C**).

**Figure 3.**
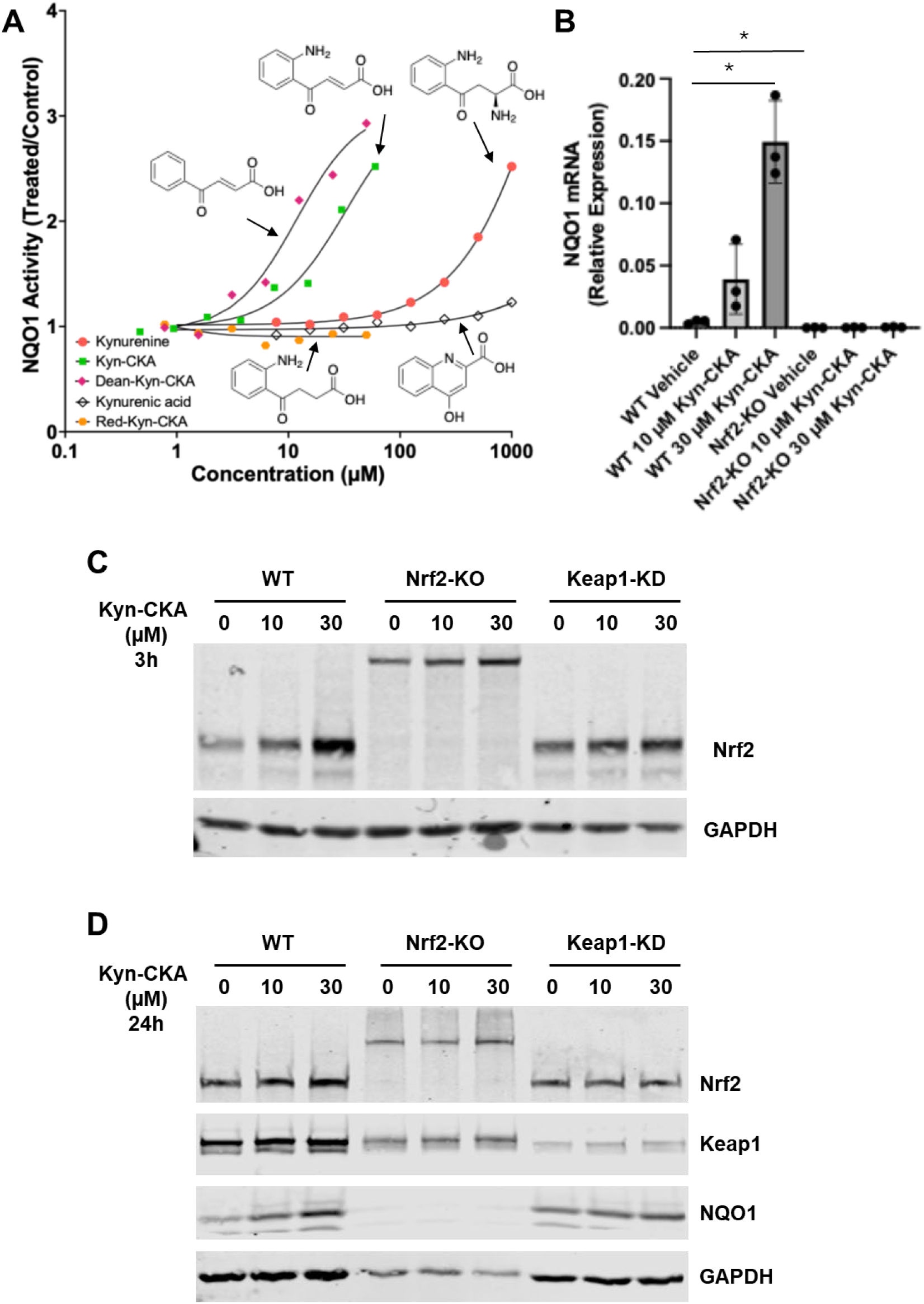
Kynurenine-CKA is a Nrf2-dependent inducer of NQO1. **(A)** Human ARPE-19 cells were seeded into 96-well plates (10,000 cells/well) and treated 24 h post-seeding with either vehicle [0.1% MeOH (v/v)] or increasing concentrations of: kynurenine, Kyn-CKA, the deaminated derivative of kynurenine Dean-Kyn-CKA, the reduced derivative of kynurenine Red-Kyn-CKA, or kynurenic acid (KynA). After 24 h, the specific enzyme activity of NQO1 was quantified in cell lysates. Data are shown as fold change in NQO1 specific enzyme activity relative to vehicle control. Values represent means of 8 biological replicates. The standard deviation was <10% of each value. Dose-response curves were fitted using a four-parameter logistic (4PL) model in GraphPad Prism. **(B)** WT and Nrf2-KO BMDMs seeded into six-well plates (1 x 10^6^/well), allowed to rest overnight, and then treated for 18 h with vehicle or Kyn-CKA (10 µM or 30 µM). The levels of mRNA for *Nqo1* were determined by quantitative RT-PCR after normalisation to the housekeeping control, *B2m*. Bars represent means ± SEM (n=3 biologically independent replicates per condition), with individual data points shown. *, p<0.05. **(C,D)** WT, Nrf2-KO and Keap1-KD BMDMs were seeded into six-well plates (1 x 10^6^/well), allowed to rest overnight, and subsequently treated with vehicle or Kyn-CKA (10 µM or 30 µM). Whole-cell lysates were collected after 3 h and 24 h and analysed by immunoblotting for Nrf2 at the 3 h timepoint **(C)** and for Nrf2, Keap1 and NQO1 at the 24h timepoint **(D)**, with GAPDH serving as a loading control.

To determine whether, similar to its kynurenine precursor, the ability of Kyn-CKA to induce NQO1 is Nrf2-dependent, BMDMs were derived from WT, Nrf2-KO and Keap1-KD mice. In unstimulated WT cells, immunoblotting showed a robust, concentration-dependent increase in Nrf2 protein levels 3 h post-treatment with Kyn-CKA (**Figure 3C**), and induction of NQO1 at the protein level at the 24 h timepoint (**Figure 3D**). In contrast, both the mRNA (**Figure 3B**) and the protein (**Figure 3D**) NQO1 levels were low, and its induction was completely abolished in Nrf2-KO BMDMs. Notably, at the 3 h timepoint, the levels of the transcriptionally inactive Nrf2-lacZ fusion protein were increased by Kyn-CKA in the Nrf2-KO BMDMs, suggesting its turnover by Keap1. Stabilisation of this fusion protein by Kyn-CKA is in agreement with previous reports that examined classical Nrf2 activators, e.g., sulforaphane in primary cultures of astrocytes obtained from this Nrf2-KO mouse strain [37]. Consistent with de-repression of Nrf2, Keap1-KD BMDMs exhibited elevated basal levels of both Nrf2 and NQO1, and Kyn-CKA treatment led to very mild induction. The protein levels of Keap1 remained largely unchanged by the Kyn-CKA treatment in each BMDM cell line (**Figure 3C**), suggesting that pathway activation occurs via post-translational modulation of Nrf2 stability rather than altered Keap1 abundance.

### 3.3. Kyn-CKA increases the thermostability of Keap1

In addition to its substrate adaptor function, which leads to Nrf2 repression, Keap1 also serves as a cellular sensor for electrophiles [38]. To investigate whether Kyn-CKA engages Keap1, we employed the cellular thermal shift assay (CETSA), a widely used method for detecting changes in the thermal stability of protein targets in response to ligand binding or conformational alterations [39]. Incubation for 1 h at 37°C of Kyn-CKA with lysates of U2OS cells stably expressing doxycycline (Dox)-inducible Keap1-mCherry fusion protein, or free mCherry as a control [28,30], increased the thermal stability of Keap1-mCherry, but not of free mCherry (**Figure 4A**), indicating that Kyn-CKA binds the Keap1 portion of the fusion protein. Next, the isothermal dose-response fingerprint CETSA (ITDRF^CETSA^) [40] was employed to quantify compound-target engagement by measuring the thermal stability of Keap1-mCherry at 64°C (the temperature yielding the greatest difference in thermal stability between the vehicle control and Kyn-CKA treatments) following incubation of cell lysates with Kyn-CKA, at a range of concentrations, for 1 h at 37°C. A sigmoidal curve was fitted to the data, from which an apparent IC_50_ value of 12 µM was calculated, representing the concentration of Kyn-CKA required for half-maximal thermal stabilisation of Keap1-mCherry (**Figure 4B**). As expected, Kyn-CKA did not alter the stability of free mCherry protein at any of the concentrations tested. Crucially, an identical ITDRF^CETSA^ experiment with Red-Kyn-CKA instead of Kyn-CKA failed to affect the thermostability of Keap1-mCherry, further confirming the importance of electrophilicity of Kyn-CKA for engaging Keap1.

**Figure 4.**
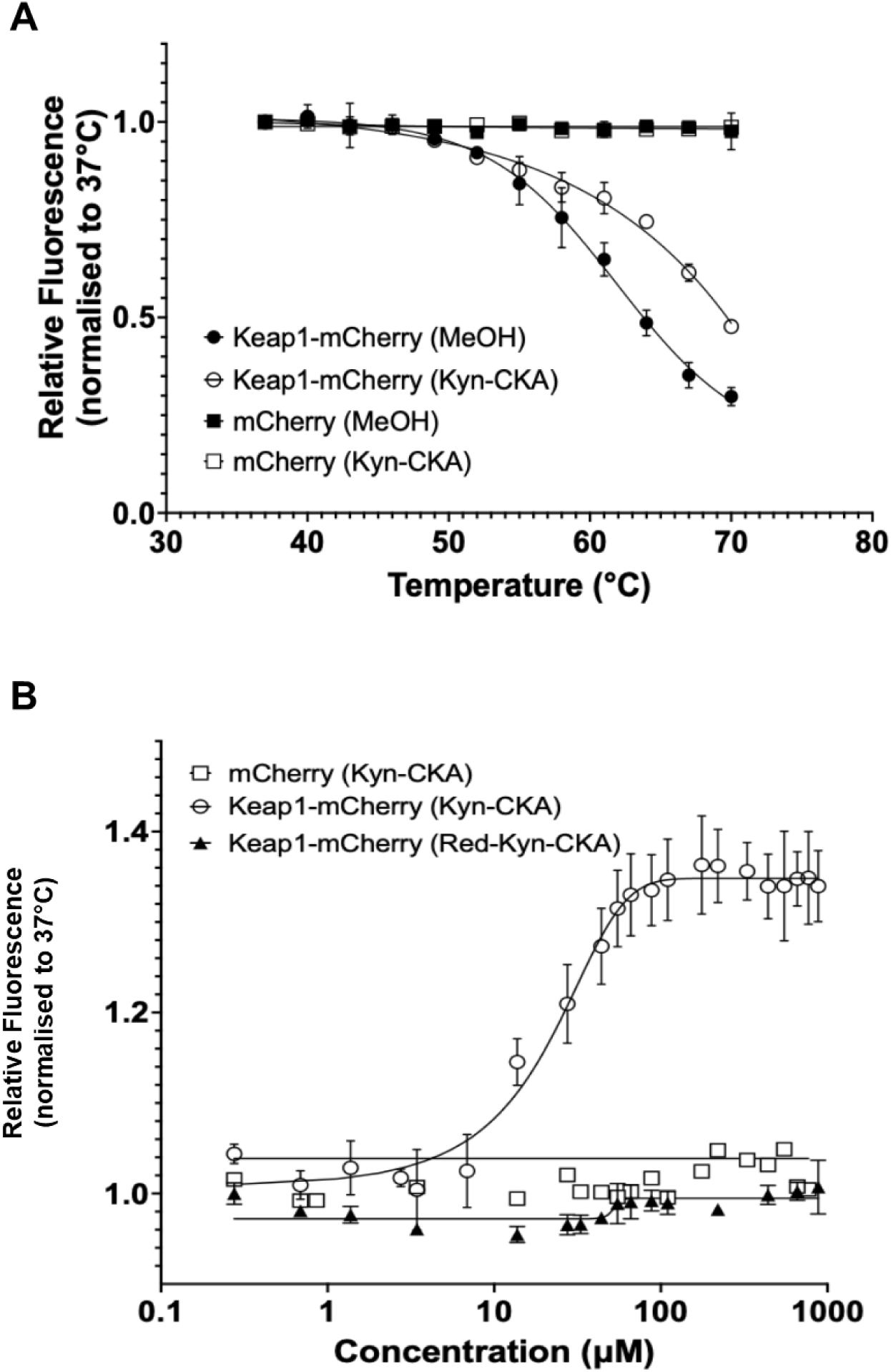
Kyn-CKA increases the thermostability of Keap1-mCherry. **(A)** Temperature-induced aggregation curves of Keap1-mCherry (n=5) and mCherry (n=3) following a 1h incubation of lysates from Keap1-mCherry- or free mCherry-expressing U2OS cells with 110 µM Kyn-CKA at 37°C. Curve fitting was performed using a four-parameter logistic (4PL) model in GraphPad Prism. **(B)** Isothermal dose-response fingerprint (ITDRF) CETSA showing concentration-dependent stabilisation of Keap1-mCherry by Kyn-CKA. Lysates of cells expressing Keap1-mCherry (n=3) or free mCherry (n=1) were incubated with Kyn-CKA for 1 h at 37°C. Lysates of Keap1-mCherry-expressing cells (n=3) were also incubated with the reduced analogue Red-Kyn-CKA. After compound incubation, cells were heated at 64°C, aggregated proteins were removed by centrifugation, and the mCherry fluorescence was measured in the soluble fraction. Data are means ± SEM. A four-parameter logistic (4PL) fit (X is log of concentration) is shown for Keap1-mCherry + Kyn-CKA; the other conditions show no concentration-dependent stabilisation.

### 3.4. C151 in Keap1 is the primary sensor for Kyn-CKA

Kyn-CKA forms covalent adducts with the reactive thiol groups of both glutathione and free cysteine [21]. Together with the observed increase in the thermostability of Keap1, this strongly suggests that Kyn-CKA can similarly modify sensor cysteine(s) within Keap1. Such covalent modification(s) would impair the substrate adaptor function of Keap1 to target Nrf2 for ubiquitination and degradation, leading to stabilisation of the transcription factor, its accumulation in the nucleus, and increased expression of Nrf2-target genes, as observed in the previous experiments (**Figure 2**). To determine the main cysteine sensor(s) of Keap1 that are responsible for the activation of Nrf2 by Kyn-CKA, we used MEF cells derived from mice expressing either wild-type (WT) Keap1 or various mutant forms of Keap1, i.e. C151S, C226S, C613S, or C288E, and the double mutant C226S/C613S [27,41]. To establish that WT MEFs respond appropriately to electrophilic stimuli comparable to the other cell models used in the study, the NQO1 enzyme activity bioassay was performed following treatment with Kyn-CKA, Dean-Kyn-CKA, and Red-Kyn-CKA.

In close agreement with the results in human ARPE-19 cells (**Figure 3A**), both Kyn-CKA (CD = 30 µM) and Dean-Kyn-CKA (CD = 12.5 µM) induced a concentration-dependent increase in NQO1 activity in WT MEFs, whereas kynurenine was a much weaker inducer, and Red-Kyn-CKA was essentially inactive (**Figure 5A**). Compared to WT cells, the mutant MEFs showed diminished inducer potency of Kyn-CKA, particularly in the C151S-Keap1 mutant MEFs (**Figure 5B**), suggesting a role for C151 in Keap1 in mediating Nrf2 activation. This conclusion was further supported by immunoblotting, which showed that treatment with 10 μM Kyn-CKA, 10 μM Dean-Kyn-CKA, or 10 nM TBE-31 (used as a positive control) increased the levels of Nrf2 in WT and all Keap1-mutant cells, except for MEFs harbouring C151S mutant Keap1 (**Figure 5C**). Concordantly, the mRNA for NQO1 was robustly induced in WT and all mutant MEFs except for the C151S-Keap1 mutant cells (**Figure 5D**), mirroring the enzyme activity data (**Figure 5A**). Interestingly, the basal *Nqo1* mRNA levels were lower in the C151S-Keap1 mutant MEFs, suggesting that this cysteine also contributes to the expression of *Nqo1* under homeostatic conditions, perhaps due to its modification by endogenously produced electrophiles during cell metabolism [16], such as the glycolytic byproduct methylglyoxal, which has been shown to engage C151 [42], or by sequestosome/p62. Experiments with purified recombinant protein containing the BTB domain of Keap1, in which C151 resides, further confirmed conjugation of Kyn-CKA to WT, but not C151S mutant, BTB-Keap1 (**Figure 6**). In addition, while Kyn-CKA required ∼40 min to react with an approximately equimolar concentration of the thiolate form of the small molecule *N*-acetyl cysteine, the same reaction with C151 in the BTB domain of Keap1 required only ∼3 min to achieve equilibrium. Thus, the residues surrounding C151 serve to specifically target Kyn-CKA to C151, catalyzing the reaction. Collectively, these results establish C151 in Keap1 as the principal sensor mediating Nrf2 activation and induction of its target genes in response to Kyn-CKA.

**Figure 5.**
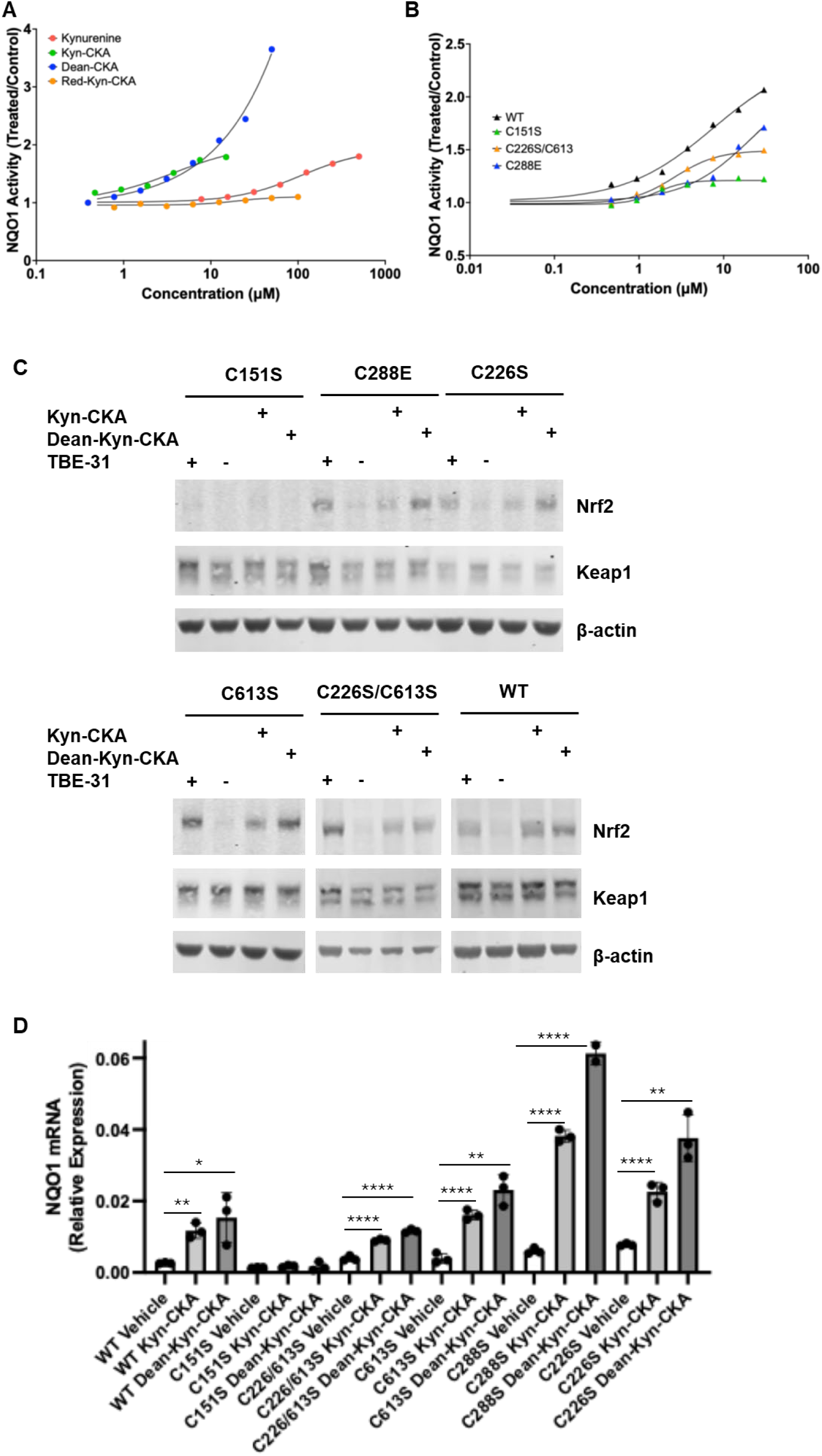
C151 in Keap1 is the primary sensor for Kyn-CKA. **(A)** MEF cells expressing wild-type Keap1 were seeded into 96-well plates (20,000 cells/well), and treated 24 h post-seeding with either vehicle [0.1% MeOH (v/v)] or increasing concentrations of kynurenine, Kyn-CKA, the deaminated derivative of kynurenine Dean-Kyn-CKA, or the reduced derivative of kynurenine Red-Kyn-CKA. After 24 h, the specific enzyme activity of NQO1 was quantified in cell lysates. Data are shown as fold change in NQO1 specific enzyme activity relative to vehicle control. Values represent means of 8 biological replicates. The standard deviation was <10% of each value. Dose-response curves were fitted using a four-parameter logistic (4PL) model in GraphPad Prism. **(B)** NQO1 enzyme activity measured using the same experimental conditions in MEF cells expressing either wild-type Keap1 (WT) or the cysteine mutants of Keap1 indicated in the figure. **(C)** WT or Keap1 mutant MEFs were treated for 3 h with 10 nM TBE-31 (a known Nrf2 activator via C151 in Keap1 used as a control), vehicle [0.1% MeOH (v/v)], 10 μM Kyn-CKA or 10 μM Dean-Kyn-CKA. Whole-cell lysates were collected and analysed by immunoblotting for Nrf2 and Keap1. β-actin served as a loading control. **(D)** Wild-type (WT) and Keap1 mutant MEFs (C151S/C151S, C288E/C288E, C226S/C226S, C226S/C613S and C613S/C613S) were treated with vehicle [0.1% MeOH (v/v)], 10 µM Kyn-CKA or 10 µM Dean-Kyn-CKA for 18 h. The mRNA levels for *Nqo1* were quantified by RT-qPCR after normalisation to the housekeeping control, *Gapdh*. Bars show means ± SEM; points denote biological replicates. *, p<0.05; **, p<0.01; ***, p<0.001; ****, p<0.0001.

**Figure 6.**
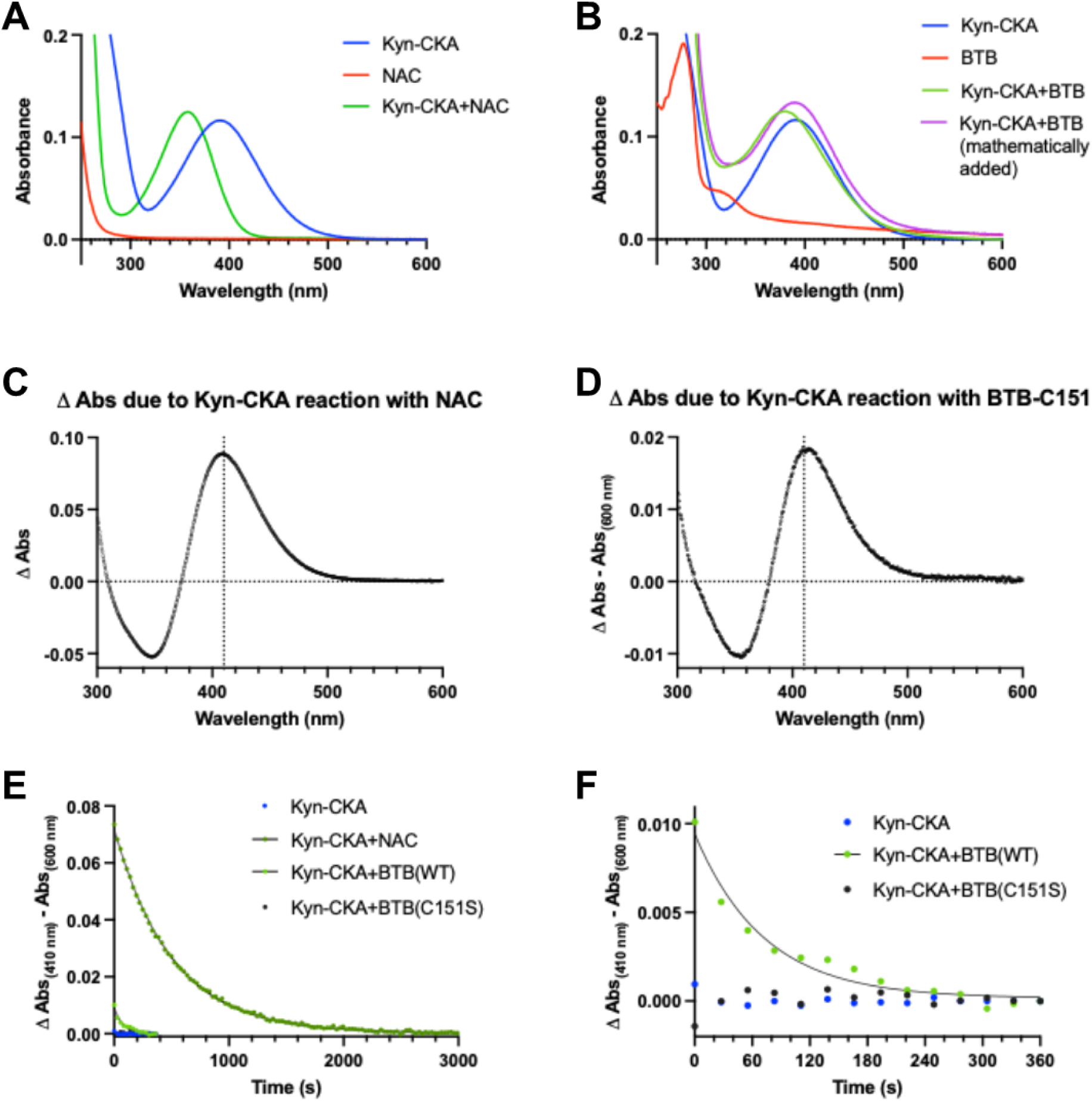
The reaction of Kyn-CKA with Keap1 C151 is catalyzed by surrounding residues in the BTB domain. **(A)** Absorbance spectra at pH 7.5 for Kyn-CKA (30 µM), *N-*acetyl cysteine (NAC) (3.5 mM, 24 µM thiolate), and the equilibrium product of their reaction. **(B)** Absorbance spectra at pH 7.5 for Kyn-CKA (30 µM), WT BTB (30 µM, 24 µM thiolate), and the equilibrium product of their reaction. The mathematical addition of the Kyn-CKA and the BTB spectra is shown in purple for comparison with the product of their reaction, in green. **(C)** The absorbance of Kyn-CKA was added to that of NAC, followed by subtraction of the product’s absorbance. The vertical dotted line marks 410 nm on the x-axis. **(D)** Calculations were done as for (C), with WT BTB absorbances in place of NAC. **(E)** The change in absorbance at 410 nm over time for the indicated reactions was plotted versus time. Data for the reactions with NAC and WT BTB were fit to a one-phase decay equation. **(F)** The data from (E) without the NAC reaction are shown graphically to better display the difference between the WT BTB and BTB-C151S proteins.

### 3.5. Kyn-CKA activates AhR, but this ligand-activated transcription factor is dispensable for the anti-inflammatory effects of Kyn-CKA

In addition to Nrf2, Kyn-CKA induces AhR-dependent gene expression while inhibiting NF-κB and NLRP3-dependent pro-inflammatory signalling [21]. The use of a mouse AhR reporter assay confirmed AhR activation by Kyn-CKA and further showed that Kyn-CKA is a more potent AhR activator than kynurenine following a 24 h incubation (**Figure 7A**). In macrophages, AhR activation suppresses NF-κB transcriptional activity resulting in decreased TNFα and IL6 production [43,44]. In agreement with these observations, the chemical inhibition of AhR with CH223191 diminished AhR activity in the reporter assay (**Figure 7B**) and resulted in a slight elevation in the mRNA levels for MCP1 and Nos2 upon BMDM challenge with LPS. However, CH223191 had no effect on the ability of Kyn-CKA to suppress the LPS-mediated pro-inflammatory responses in these cells (**Figure 7C-F**). In agreement with the mRNA data, AhR inhibition did not affect the ability of Kyn-CKA to decrease the secreted levels of MCP1 and IL6 in the cell culture media (**Figure 7G,H**).

**Figure 7.**
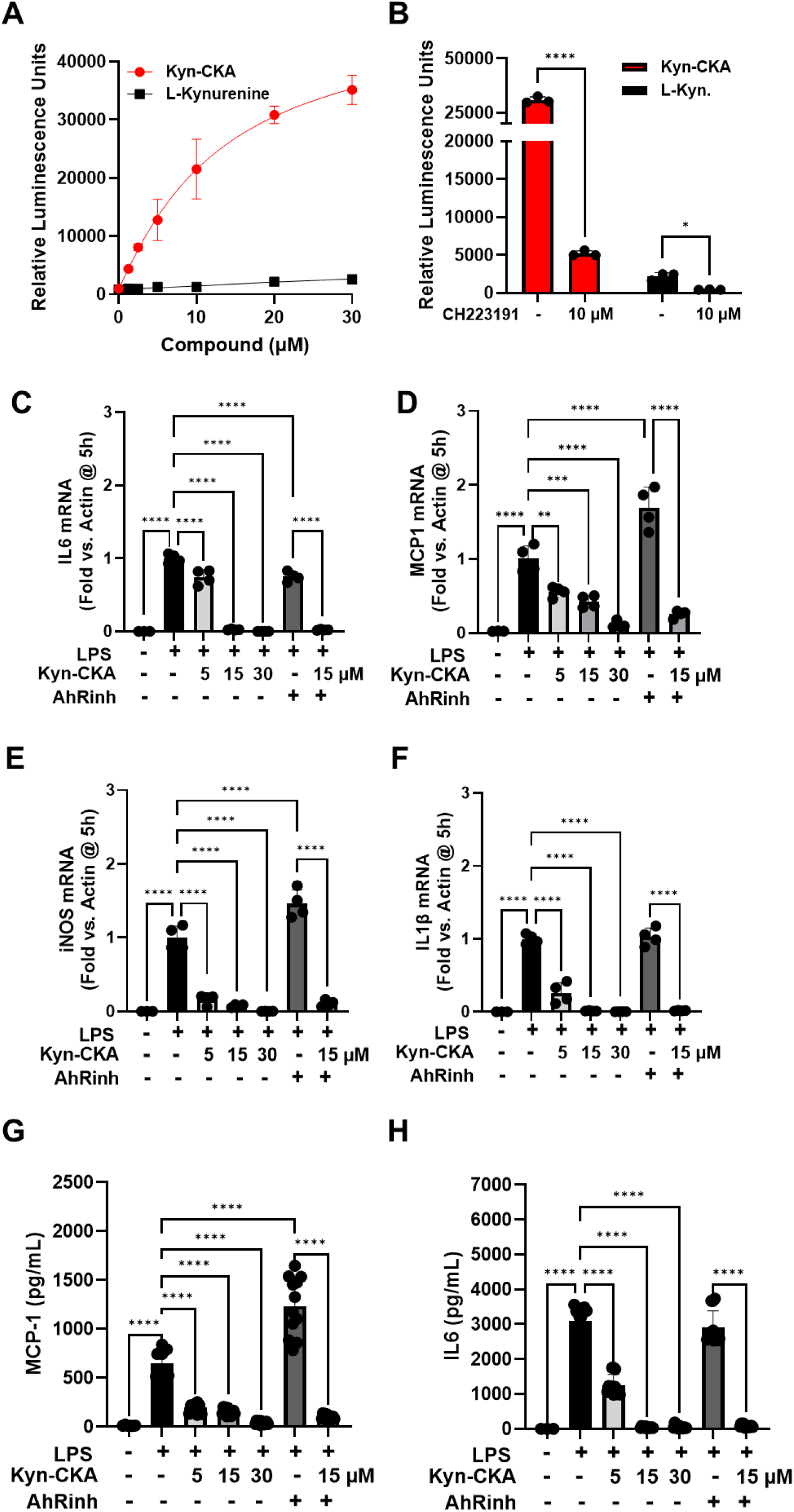
AhR inhibition has no effect on the anti-inflammatory actions of Kyn-CKA in BMDMs. **(A)** Mouse AhR reporter cells expressing luciferase under the control of the xenobiotic response element (XRE) were incubated with Kyn-CKA or kynurenine for 24 h prior to luminescence measurement (n=3). Data were fitted to a sigmoidal four-parameter logistic curve (L-Kyn: r^2^ = 0.824, IC_50_ = 28 µM, span = 3,218 RFU. Kyn-CKA: r^2^ = 0.970, IC_50_ = 13 µM, span = 47,901 RFU). **(B)** Inhibition of AhR-dependent luciferase expression by CH223191 (AhRinh). **(C-F)** Kyn-CKA inhibits the expression of the NF-κB-regulated genes IL6, MCP1, Nos2 and IL1β in BMDM following 5 h incubation with LPS (0.5 µg/mL) and the indicated concentrations of Kyn-CKA both in the presence or absence of CH223191 (10 µM). Data are n=3-4, **** p<0.0001 by one-way ANOVA and Tukey’s post-test. **(G-H)** Extracellular cytokine levels from BMDM treated as in C-F for 5 h. Data are n=3-4 (all replicates shown), **** p<0.0001 by one-way ANOVA and Tukey’s post-test.

Moreover, the deaminated derivative of Kyn-CKA, DEAN-CKA, which activated Nrf2 (**Figure 3A**), but did not activate AhR (**Figure 8A**), also lowered the levels of secreted IL6 and MCP1 from LPS-stimulated BMDMs (**Figure 8B,C**), further indicating that activation of AhR is not required for the anti-inflammatory activity of compounds of this type.

**Figure 8.**
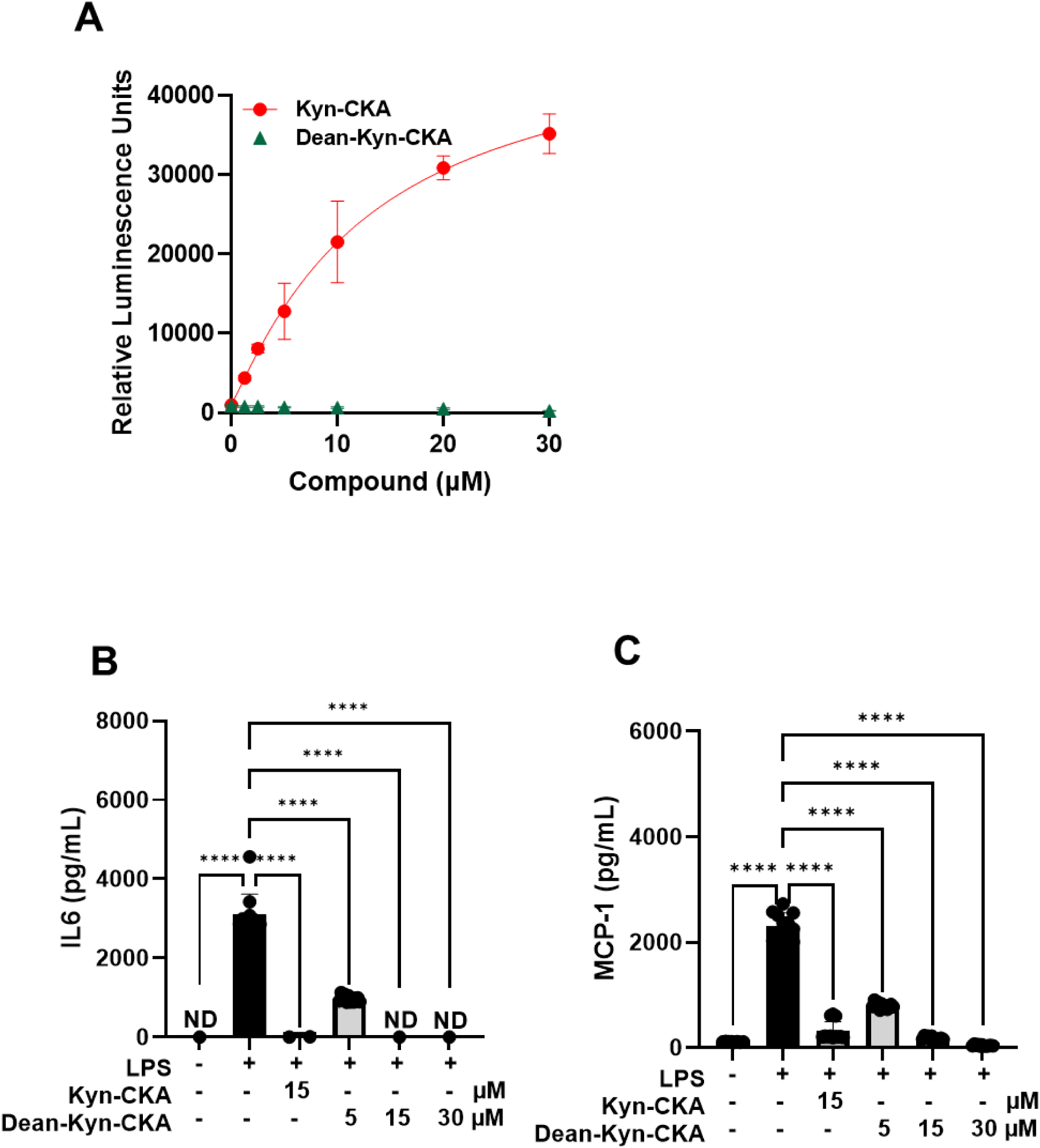
The AhR-inactive DEAN-CKA inhibits LPS-induced pro-inflammatory gene expression with similar potency to Kyn-CKA. **(A)** Mouse AhR reporter cells expressing luciferase under control of the xenobiotic response element (XRE) were incubated with Dean-Kyn-CKA or Kyn-CKA for 24 h prior to luminescence measurement (n=3). Data were fitted as in figure 7 (Dean-Kyn-CKA: No fit. Kyn-CKA: r^2^ = 0.970, IC_50_ = 13 µM, span = 47,901 RFU). **(B-C)** Extracellular cytokine levels from BMDM treated as in C-F for 5 h. Data are n=3-4, **** p<0.0001 by one-way ANOVA and Tukey’s post-test.

Experiments in BMDMs derived from monocyte-specific AhR-knockout (*LysM Ahr^-/-^*) mice confirmed that Kyn-CKA treatment leads to transcriptional upregulation of Nrf2-target genes (**Figure 9A-C**), indicating that the absence of the AhR does not affect the ability of Kyn-CKA to activate Nrf2. Most importantly, these experiments further showed that the downregulation of the LPS-mediated transcription of pro-inflammatory genes by Kyn-CKA does not require the presence of the AhR (**Figure 9D-I**). Together, these data establish firmly that the AhR is dispensable for the anti-inflammatory activity of Kyn-CKA in macrophages.

**Figure 9.**
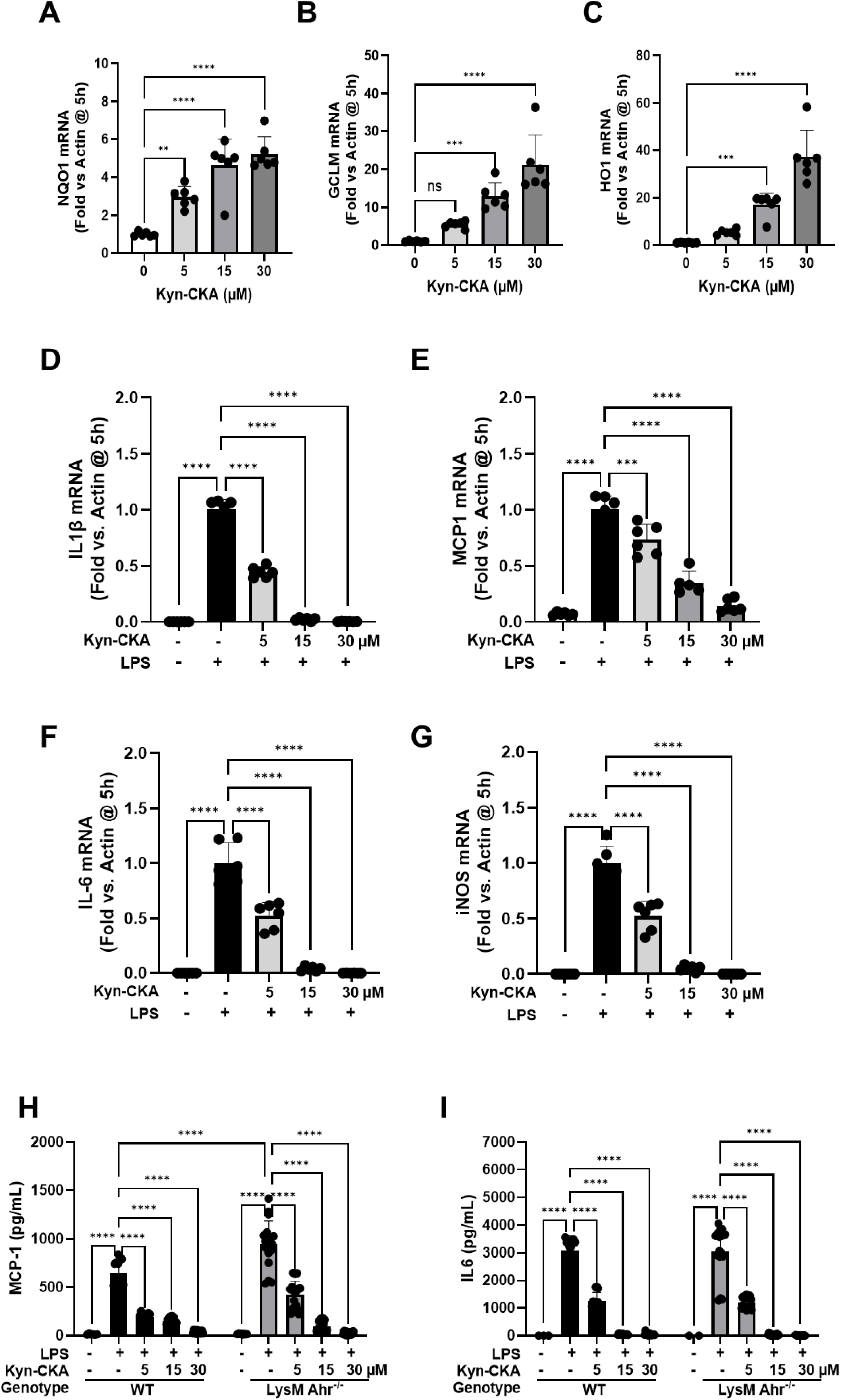
Kyn-CKA inhibits LPS-elicited responses in AhR-knockout BMDMs. **(A-C)** Kyn-CKA dose-dependently induces Nrf2-dependent gene expression in BMDM obtained from LysM AhR-knockout (AhR^-/-^) mice. Data are n=6, ** p<0.01, *** p<0.001, **** p<0.0001 by one-way ANOVA and Dunnett’s post-test. **(D-G)** Kyn-CKA inhibits the expression of the NF-κB-regulated genes in LysM AhR^-/-^ BMDM following 5h incubation with LPS (0.5 µg/mL) and the indicated concentrations of Kyn-CKA. Data are n=6, **** p<0.0001 by one-way ANOVA and Dunnett’s post-test. **(H-I)** Extracellular cytokine levels from WT and LysM AhR^-/-^ BMDM treated as in D-G for 5 h. Data are n=6, **** p<0.0001 by two-way ANOVA and Tukey’s post-test. The WT BMDM data were replotted from figure 7G-H for comparison purposes.

### 3.6. The ability of Kyn-CKA to suppress pro-inflammatory responses is partly Nrf2-dependent

To test whether Nrf2 contributes to the anti-inflammatory activity of Kyn-CKA, we used BMDMs from WT and Nrf2-KO mice. Nrf2 activation by Kyn-CKA was confirmed by the increase in mRNA levels for Nqo1 and Gclm, which occurred in WT, but not Nrf2-KO cells (**Figure 10A,B**). As expected, Kyn-CKA treatment of WT BMDMs, led to a concentration-dependent downregulation of the LPS-stimulated transcription of the pro-inflammatory markers MCP1, IL1β, IL6, TNFα and Nos2 (**Figure 10C-G**). In contrast, the suppressive effect of the low (5 µM) concentration of Kyn-CKA was not apparent in Nrf2-KO BMDMs, but was still observed when higher Kyn-CKA concentrations were used; this effect was particularly evident at the highest (30 µM) concentration of Kyn-CKA. Together, these experiments suggest that, at low concentrations, Keap1/Nrf2 mediate the anti-inflammatory effect of Kyn-CKA, but at high concentrations, Kyn-CKA affects other proteins that are involved in the inflammatory response.

**Figure 10.**
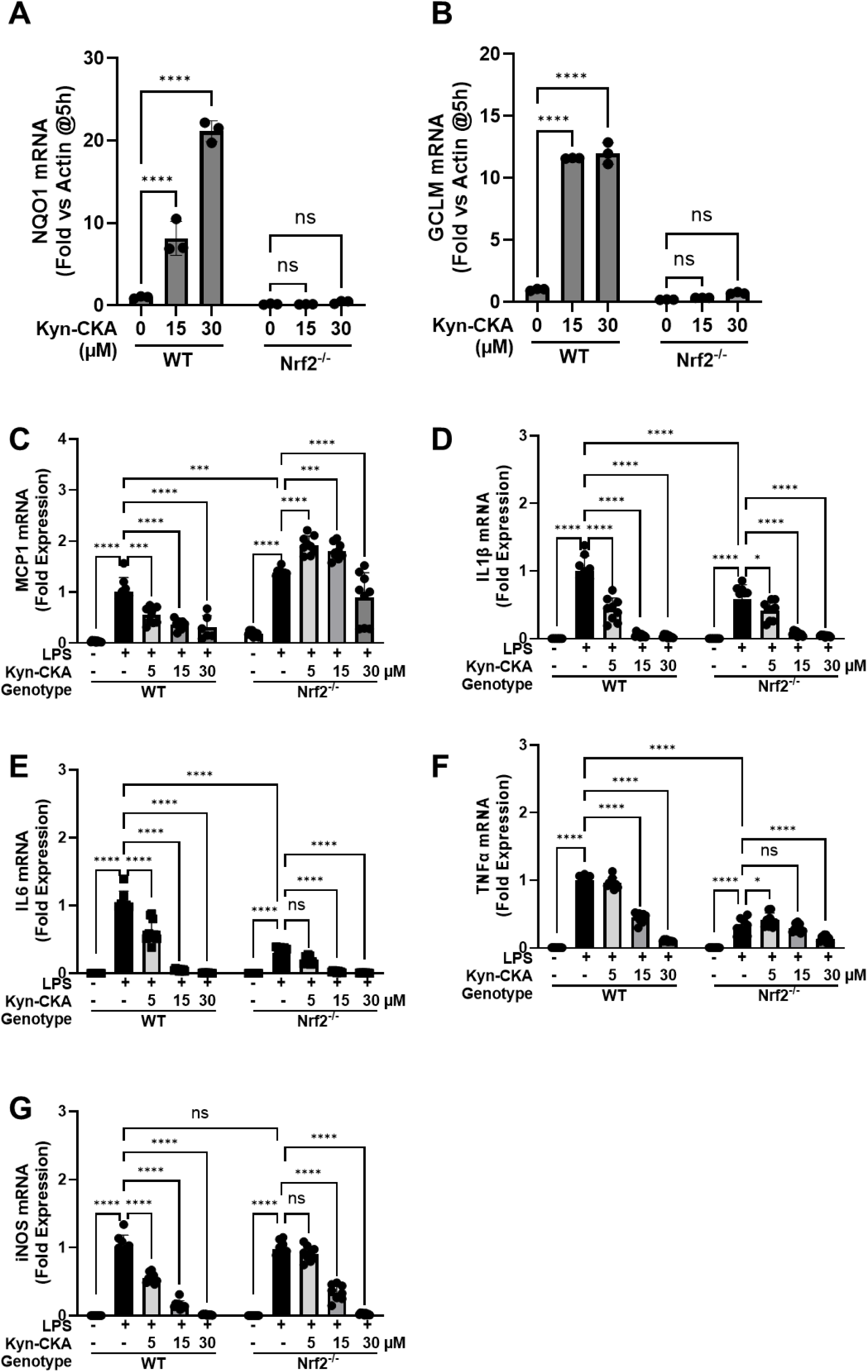
The low-dose anti-inflammatory effects of Kyn-CKA in BMDMs are dependent on Nrf2. **(A-B)** Treatment with Kyn-CKA (5 h) fails to induce the expression of *Nqo1* and *Gclm* in BMDMs obtained from Nrf2-knockout (Nrf2^-/-^) mice. Data are n=3, **** p<0.0001 by two-way ANOVA and Tukey’s post-test. **(C-G)** Effects of Kyn-CKA on the expression of NF-κB−regulated genes in WT and Nrf2-KO BMDMs following 5 h incubation with LPS (0.5 µg/mL) and the indicated concentrations of Kyn-CKA. Data are n=6, * p<0.05, *** p<0.001, **** p<0.0001 by two-way ANOVA and Tukey’s post-test. Nrf2 was identified as a significant source of variation for MCP1, IL6, TNFα, Nos2, (p<0.0001) and IL1β (p<0.005).

## 4. Discussion

Keap1 was discovered ∼25 years ago as the principal negative regulator of Nrf2 [45]. Shortly afterwards, it was demonstrated that Keap1 is endowed with reactive cysteines that function as molecular sensors for a wide variety of small molecules, most of them xenobiotics, all of which have sulfhydryl reactivity as their common chemical property [46]. More recently, several endogenous electrophiles have been shown to modify cysteines in Keap1; these have been reviewed [16]. Such modifications inactivate the substrate adaptor activity of Keap1, resulting in de-repression of Nrf2 and induction of a broad cytoprotective program that allows adaptation and survival under conditions of cellular stress, restoring homeostasis. One putative endogenous Nrf2 activator is the tryptophan metabolite kynurenine [33]. However, kynurenine lacks electrophilic reactivity, and the mechanism by which it activates Nrf2 was not known hitherto. Here, we confirm that kynurenine is a low-potency (high micromolar range) activator of Nrf2. We further show that its electrophilic metabolite Kyn-CKA is more potent than its kynurenine precursor and induces Nrf2-mediated transcription through reacting with C151 within Keap1. Together with our previous report that kynurenine synthesis in cells, such as macrophages, leads to Kyn-CKA formation and that Kyn-CKA–specific metabolites have been detected in humans and mice [21], the findings from the current study strongly suggest that Kyn-CKA rather than kynurenine, is the *de facto* inducer.

In macrophages, genetic or pharmacological Nrf2 activation, including by Kyn-CKA, suppresses pro-inflammatory responses [21]. In addition to Nrf2, kynurenine and Kyn-CKA promote AhR-dependent gene expression [21]. Activation of AhR has anti-inflammatory effects; and this has been shown in animal models of human disease, such as autoimmune encephalomyelitis and retinal ischemia/reperfusion injury [47,48]. Thus, in this study we considered the possibility that the anti-inflammatory effects of Kyn-CKA might be mediated, at least in part, by AhR. The use of BMDMs derived from monocyte-specific AhR-knockout mice revealed that, although Kyn-CKA robustly activates AhR in these cells, the anti-inflammatory effect of this metabolite is AhR-independent. In contrast, our experiments in BMDMs derived from Nrf2-KO mice showed a clear Nrf2-dependence at low Kyn-CKA concentrations that likely reflect its endogenous cellular levels [21].

Interestingly, in LPS-stimulated BMDM cells, the expression of IL1β, IL6 and TNFα was lower in the absence of Nrf2. This finding is in close agreement with our earlier results in SKH1- hairless mice exposed to solar-simulated ultraviolet radiation (SSUV) showing that, whereas Nrf2 is required for the anti-inflammatory effect of TBE-31, the SSUV-induced expression of IL1β and IL6 in the skin of Nrf2-KO mice was lower than in the skin of their wild-type counterparts [24]. A recent RNA-seq analysis has revealed that Nrf2 deficiency in microglia dampens the microglial pro-inflammatory responses to systemic LPS administration [49]. Further investigations are necessary to elucidate the underlying mechanism(s), but these observations strongly suggest that, in addition to its widely recognized role in the resolution of inflammation, Nrf2 might also support the inflammatory response.

Notably, at high concentrations, Kyn-CKA was still able to suppress the transcription of pro-inflammatory genes in the absence of Nrf2. These results are consistent with our previous findings in murine peritoneal macrophages stimulated with IFNγ and TNFα, where two potent Nrf2 activators (SFN and the pentacyclic cyanoenone TP-225) were much less effective inhibitors of Nos2 induction in Nrf2-KO in comparison with WT cells, but inhibition still occurred when higher concentrations of these electrophiles were used [15]. We suggest that the Nrf2-independent effects of Kyn-CKA are mediated by inhibition of NF-κB signalling potentially associated with direct covalent modification of p65 [21], but that the Keap1/Nrf2 system, which is affected by the low concentrations of Kyn-CKA, is the primary target. This scenario is reminiscent of the ‘‘hierarchical’’ or ‘‘rheostat’’ response of the murine macrophage cell line RAW 264.7 to diesel exhaust particles, which contain redox cycling organic chemicals that induce pro-oxidative and pro-inflammatory effects [50]. In that response, the Keap1/Nrf2 system is the first-tier defence (activated by modest increases of reactive oxygen / nitrogen species (ROS/RNS), whereas NF-κB represents the second-tier defence (activated by higher ROS/RNS levels); the third and final tier involves activation of apoptosis. Thus, as described for conditions of oxidative stress [50], the Keap1/Nrf2 system may serve as a ‘‘floodgate’’ for electrophiles, such as Kyn-CKA, in which other transcription factors are affected at a particular threshold of concentration only after “saturation” of the Keap1 cysteine sensor(s).

## Acknowledgments

We thank the staff at the WTB Resource Unit of the University of Dundee for technical assistance, the Ninewells Cancer Campaign Doctoral Training Programme, and the Medical Research Council (MR/W023806/1 and MR/T014644/1) for funding our research. We would also like to acknowledge support by National Institutes of Health grants R35GM152083 (DAV), R01AI178864 (DAV) and T32AI007051 (NAD), as well by a grant from the College of Liberal Arts and Sciences at Villanova University.

## CRediT authorship contribution statement

Jialin Feng: Investigation, Data curation and analysis, Writing – review & editing. Mara Carreño: Investigation, Data curation and analysis. Hannah Jung: Investigation, Data curation and analysis. Sharadha Dayalan Naidu: Methodology, Supervision. Nicole Arroyo-Diaz: Methodology, Investigation. Abel D. Ang: Resources. Bhargavi Kulkarni: Investigation. Dorothy Kisielewski: Resources. Takafumi Suzuki: Resources, Writing – review & editing. Masayuki Yamamoto: Writing – review & editing. John D. Hayes: Writing – review & editing. Tadashi Honda: Resources. Beatriz Leon-Ruiz: Conceptualization, Resources. Aimee L. Eggler: Conceptualization, Funding acquisition, Supervision, Data curation, Writing – review & editing. Dario A. Vitturi: Conceptualization, Funding acquisition, Supervision, Data curation, Writing – review & editing. Albena T. Dinkova-Kostova: Conceptualization, Funding acquisition, Supervision, Data curation, Writing – original draft, review & editing.

